# The Auxin and Cytokinin Balance Influence the in Vitro Regeneration of *Phalaenopsis* Shoots (Orchidaceae)

**DOI:** 10.1101/2023.07.11.548545

**Authors:** Kaliane Zaira Camacho Maximiano Cruz, Antonio André Silva Alencar, Josefa Grasiela Silva Santana, Laura Eliza Oliveira Alves

## Abstract

Orchids (*Phalaenopsis*) are considerably appreciated or their high and durable flowering rate and exotic appearance. The optimization of in vitro cultivation begins with an understanding of hormonal balance and its implications in production. This work aimed to regenerate shoots from seeds germination of the *Phalaenopsis* Golden Peoker ‘BL’ HCC / AOS’ under exogenous auxin and cytokinin influence. To do so, naphthalene acetic acid (NAA), 6-benzyladenine (BA), and Kinetin (Kin) were added to the culture medium to induce organogenesis, totaling 17 treatments with eight replications per treatment. The variables number of shoots (NS), shoot length (SL), and presence of root primordia (RP) were evaluated. A non-parametric test (the Friedman test), followed by the Bonferroni procedure, was used to compare the different groups. Hence, the MS medium provided 610 complete plants after 90 days of cultivation. The T6 treatment had the highest median NS (1.5 shoots per plant at 15 days) and SL. At 30 days, the highest NS was for the T16, with 6.15 shoots per plant; for SL, the one that stood out was the T9. Regarding RP in shoots, the best treatments were T2, T3, T9, T10, and T17, with the formation of RP at 15 days, remaining these treatments at 30 days. In conclusion, the T9 is the most suitable for an efficient protocol reproduction, as it showed the best plant development.

**HIGHLIGHTS (MANDATORY):**

- The auxin and cytokinin balance promoted shoot regeneration.
- The BA and NAA combined promoted the shoot formation of *Phalaenopsis* Golden Peoker ‘BL’ HCC/AOS’.
- The use of KIN isolated showed hyperhydricity and malformation in shoots of *Phalaenopsis*.

## INTRODUCTION

Among the most cultivated orchids, the *Phalaenopsis*—originally from Southeast Asia—stands out for its appearance, showing exotic flowers, a high floral count, and the durability of its inflorescences [1].

Due to their use in cultural practices [2], they are economically significant ornamental cut flowers and potted floricultural crops worldwide [2, 3].

Developing new varieties and hybrids to grow beautiful new plants is considered one of the main factors in the commercial demand for flowers and ornamental plants, which has increased as consumer preferences and needs have grown [4].

Hundreds of hybrids have been introduced to the world market over the last decades, and, bearing in mind the seedling production system, there has been massive investment in various countries, such as Thailand, Taiwan, Vietnam, and China, where the production of millions of seedlings is done through asymbiotics carried out in industrial laboratories [4]. Because of its growing demand and commercial investment, more studies and research about orchid propagation techniques have become relevant.

In the orchid plant tissue culture, besides seed germination in vitro, other morphogenetic routes can be investigated, such as in vitro regeneration or morphogenesis. This route leads to the organ development from other pre-existing organs and may occur by the embryogenesis or organogenesis route [5].

In vitro organogenesis is a process that involves several phases, such as dedifferentiation; acquisition of competence; induction; determination; differentiation and regeneration or formation of an organ [6]. Therefore, this process becomes a viable alternative for clonal propagation since, from the mother plant, the shoot formation that later develops into plants may occur extensively.

Throughout the process of organogenesis, the auxin/cytokinin balance represents a critical factor in plant growth regulation [7]. Auxin and cytokinin have antagonistic effects on cell division and differentiation, with auxin promoting cell division, and cytokinin, cell differentiation. Thus, the ratio of auxin to cytokinin is crucial to determining the fate of cells and the type of tissue formed. Understanding the interplay between these hormones is essential for optimizing tissue culture protocols and developing new strategies for plant regeneration and propagation [8].

Using plant micropropagation from in vitro morphogenesis has been a reality in several countries—for example, Brazil— and is already being applied to large-scale commercial production of the genus *Phalaenopsis* [9]. The aim of this work was to regenerate shoots from seeds of the *Phalaenopsis* Golden Peoker ‘BL’ HCC / AOS’ under the influence of exogenous auxin and cytokinin.

## MATERIAL AND METHODS

### Asepsis and in vitro germination

A hybrid orchid capsule of *Phalaenopsis* Golden Peoker ‘BL’ HCC / AOS’ was disinfected in a laminar flow chamber under aseptic conditions through immersion in 70% ethanol (Sigma – Aldrich, St. Louis, USA) for 1 min, followed by immersion in 2.5% commercial bleach (sodium hypochlorite from 0.6 to 0.75%; Qboa® Anhembi AS, Osasco, Brazil) supplemented with two drops of Tween® 20 (Sigma – Aldrich) for 30 min, and subsequently by four rinses with distilled and autoclaved water.

Inside a laminar flow chamber, the seeds were aseptically removed from the capsule and transferred into a glass flask (50 × 100 mm) containing 50 mL of MS [10] culture medium supplemented with 30 g L^-1^ of sucrose, 100 mg L^-1^ of myo-inositol, and solidified with 8.0 g L^-1^ of bacteriological agar (Sigma – Aldrich).

The pH was adjusted to 5.7 before autoclaving (for 15 min at 121 ºC and 1.1 atm pressure). The flasks with the culture medium were sealed using PVC film (polyvinyl chloride) – Rolopac® and maintained in a greenhouse at a light photoperiod of 16:8 h, with an irradiance of 36 μmol m^-2^ s^-1^ and temperature of 25 ± 2 ºC for in vitro seed germination for 90 days.

### Organogenesis induction

After 90 days of in vitro cultivation in a germination medium, all plants were transferred to flasks with the MS culture medium containing 30 g L^-1^ of sucrose, 100 mg L^-1^ of myo-inositol, solidified with 8.0 g L^-1^ of bacteriological agar (Sigma–Aldrich), at pH 5.7. For inducing organogenesis, different concentrations and combinations of the growth regulator, naphthalene acetic acid (NAA), 6-benzyladenine (BA), and Kinetin (KIN) were added to the culture medium, totaling 17 treatments with eight replications per treatment and each plant being one repetition (Table 1).

**Table 1:** Treatments used to induce organogenesis in *Phalaenopsis* sp. after 90 days of *in vitro* culture (seedling medium)

| Treatment | Concentration mg |
| --- | --- |
| T1 | MS0 |
| T2 | 1.0 mg BA |
| T3 | 1.5 mg BA |
| T4 | 2.0 mg BA |
| T5 | 2.5 mg BA |
| T6 | 1.0 mg BA + 0.5 mg NAA |
| T7 | 1.5 mg BA + 0.5 mg NAA |
| T8 | 2.0 mg BA + 0.5 mg NAA |
| T9 | 2.5 mg BA + 0.5 mg NAA |
| T10 | 1.0 mg KIN |
| T11 | 1.5 mg KIN |
| T12 | 2.0 mg KIN |
| T13 | 2.5 mg KIN |
| T14 | 1.0 mg KIN + 0.5 mg NAA |
| T15 | 1.5 mg KIN + 0.5 mg NAA |
| T16 | 2.0 mg KIN + 0.5 mg NAA |
| T17 | 2.5 mg KIN + 0.5 mg NAA |

The plants were grown in flasks sealed with PVC (polyvinyl chloride) film (Rolopac®) and maintained in a growth room under irradiance of 36 mmol m^-2^ s^-1^, light photoperiod of 12 h, and temperature of 25 ± 2 ºC.

After 90 days of in vitro cultivation in the medium to promote organogenesis, plants measuring approximately 5 cm were washed in distilled water to remove residues from the culture medium. Subsequently, they were acclimatized in plastic cups containing sphagnum substrate moistened with 5 mL of hypochlorite of sodium at a concentration of 0.5%. They were kept in a greenhouse at room temperature and at 70% light. The irrigations were maintained through automatic irrigations four times a day daily.

### Statistics

The experiments were carried out following a completely randomized design. In organogenesis, the variables number of shoots (NS), shoot length (SL), and presence of root primordia (RP) were evaluated. The organogenesis evaluations occurred at 15 and 30 days of in vitro culture. The median of the data set is depicted in graphs. Given the lack of normality of the errors, the Friedman test a non parametric test free of distribution was used to compare the various groups, followed by the Bonferroni procedure. A *ρ* value less than 0.05 was regarded as significant.

## RESULTS AND DISCUSSION

### Seed germination of *Phalaenopsis* Golden Peoker ‘BL’ HCC / AOS’

It was possible to observe the seed germination of the hybrid *Phalaenopsis* Golden Peoker ‘BL’ HCC/AOS’ (Figs. 1A and 1B) efficiently under aseptic conditions and after being inoculated in MS medium (10); however, the exact initial number of orchid seeds in a capsule is unknown, as they can vary from thousands to millions per capsule. The development of the protocorms initially occurred at 30 days of in vitro culture in the germination medium (basic salts of MS) (Fig. 1C).

**Figure 1.**
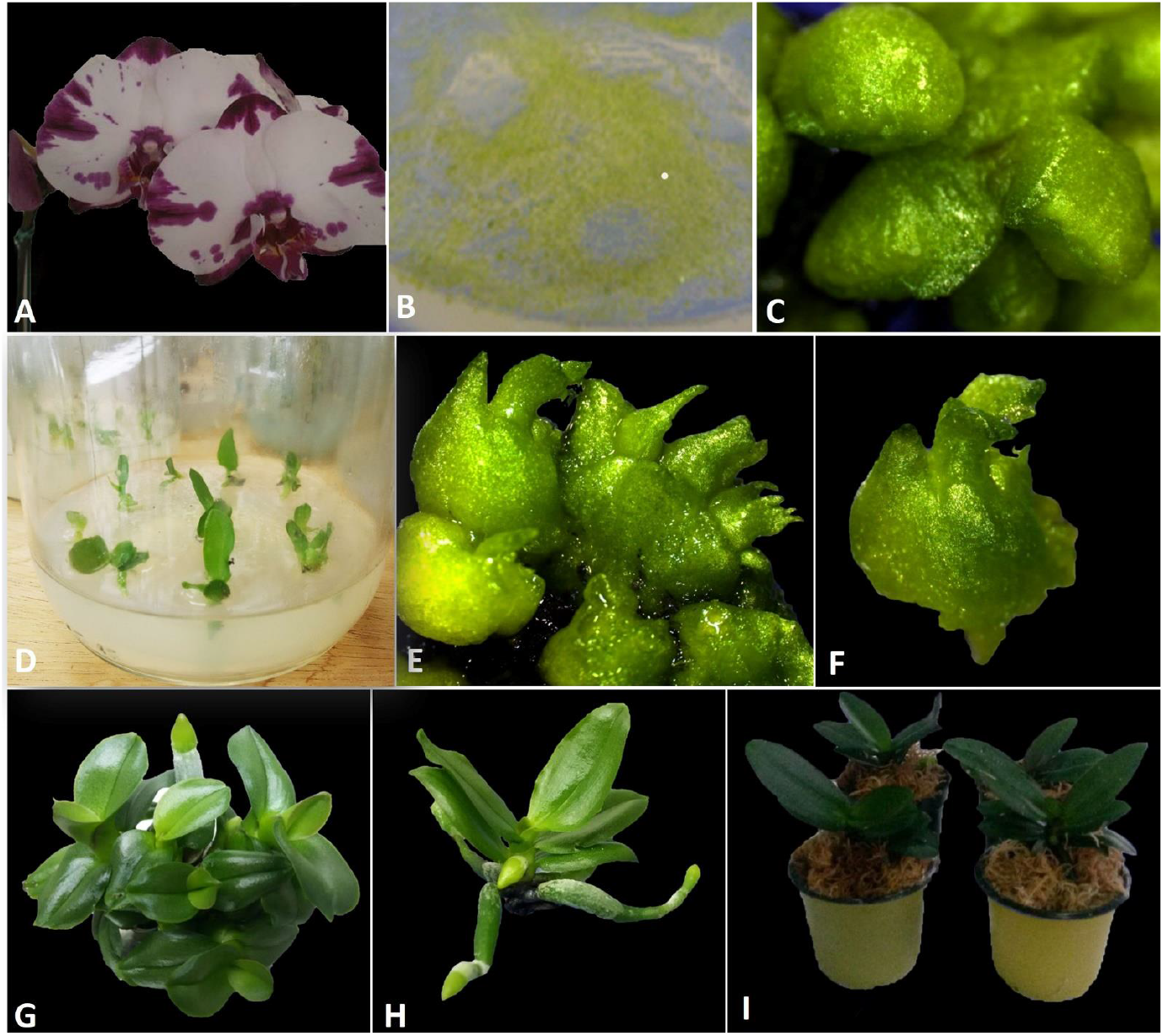
In vitro cultivation and general aspects of shoot development from direct organogenesis of *Phalaenopsis* Golden Peoker ‘BL’ HCC/AOS’. **A)** Flower of *P*. Golden Peoker. **B)** Seeds inoculated in vitro with MS medium culture. **C)** Protocorms formed after 30 days of in vitro culture. **D)** Plants with 2 cm used for organogenesis induction. **E)** Shoots formed in the T13 (2.5 mg Kin) treatment showed signs of hyperhydricity. **F)** Isolated shoot from T13 (2.5 mg Kin) treatment showed malformation and abnormal development after 30 days of culture. **G)** Shoots developed in the T9 (2.5 BA + 0.5 mg NAA) showed normal development (elongation, expanded leaves, and roots). **(H)** Isolated shoot from T9 (2.5 mg BA + 0.5 mg NAA) treatment after 30 days for acclimatization. **(I)** Plants in an acclimatized stage in sphagnum substrate with normal development in *ex vitro* process.

Unlike angiosperms in general, orchid seeds show a pattern of germination and initial growth with the embryo swelling at the beginning, which causes the seed testa to rupture and then release [11]. Similar results were observed by Koene et al., [12] using MS medium in the asymbiotic germination of *Acianthera prolifera* during the first two weeks, with the same pattern of testa rupture and germinated seeds.

The MS culture medium is widely used because it has a high concentration of salts, such as nitrogen and potassium. According to Kauth, Dutra (13), nitrogen plays a relevant role in orchid seed germination. The best initial seed germination responses in MS medium may be attributed to the higher nitrogen concentration when compared to other cultivation media. Thus, the proper choice of cultivation medium has a crucial influence on the initial stage of orchid development. After this phase, the embryo develops into a tuberiform, chlorophyll structure, the protocorm [14].

After forming the protocorms, at 60 days of the in vitro cultivation, the development of leaf primordia occurred, and, at 90 days of cultivation, 610 complete plants were recorded (Fig. 1 D) within the sowing period.

### Effects of cytokinin on organogenesis and in vitro plant growth

All treatments exhibited direct organogenesis. As the data were not normally distributed, a Friedman test (using a median) was carried out, and the frequency data for NS was shown as a supplementary figure. At 15 and 30 days of cultivation in organogenesis medium, it was found that the treatment T13, with 2.5 mg Kin, showed the highest median for shoot production, with three shoots per plant at 15 days (Fig. 2) and seven shoots at 30 days (Fig. 2). However, it was seen that the regenerants did not develop but presented an appearance of hyperhydricity (Figs. 1E and 1F), with abnormal formation during the organogenesis process.

**Figure 2.**
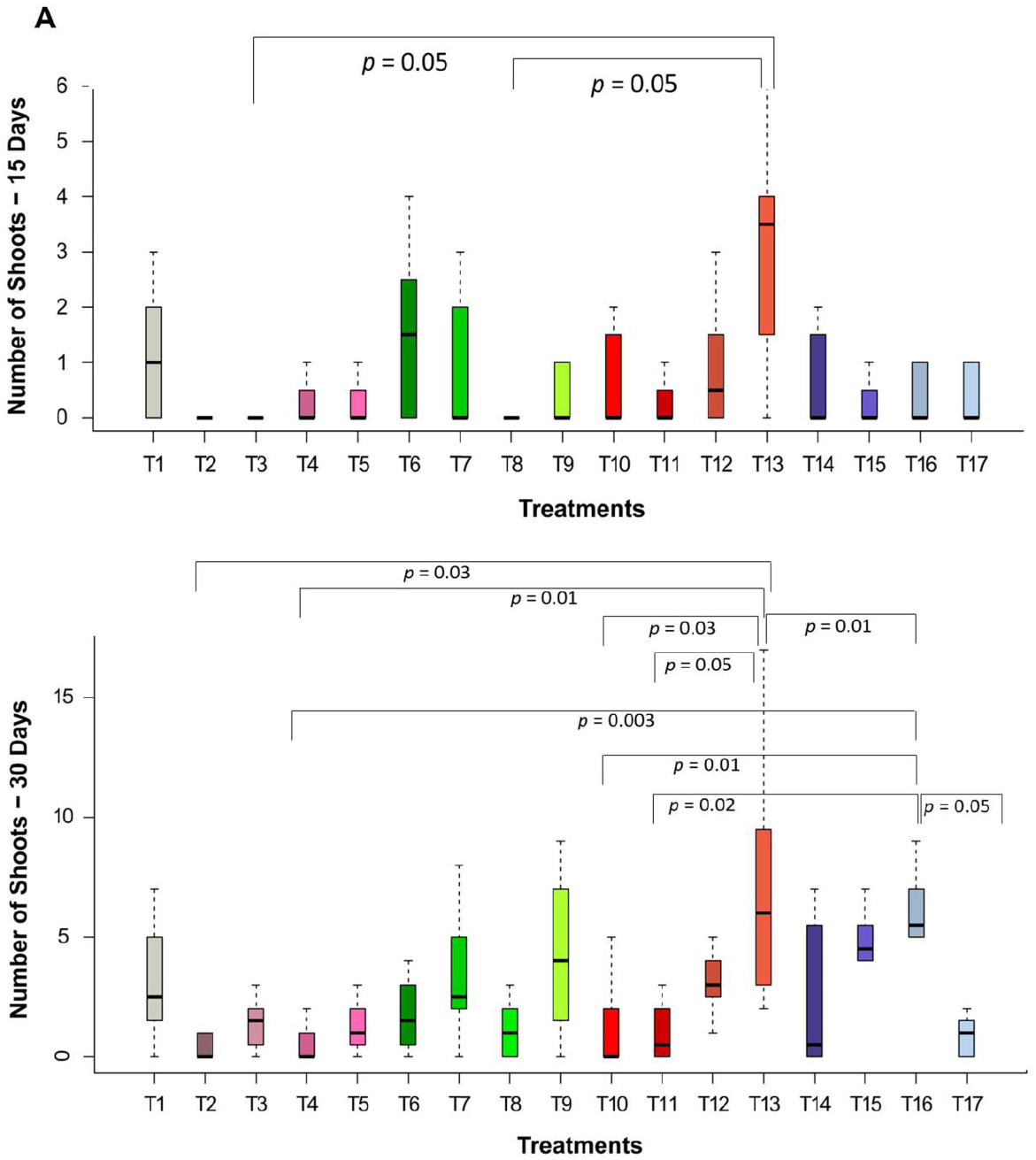
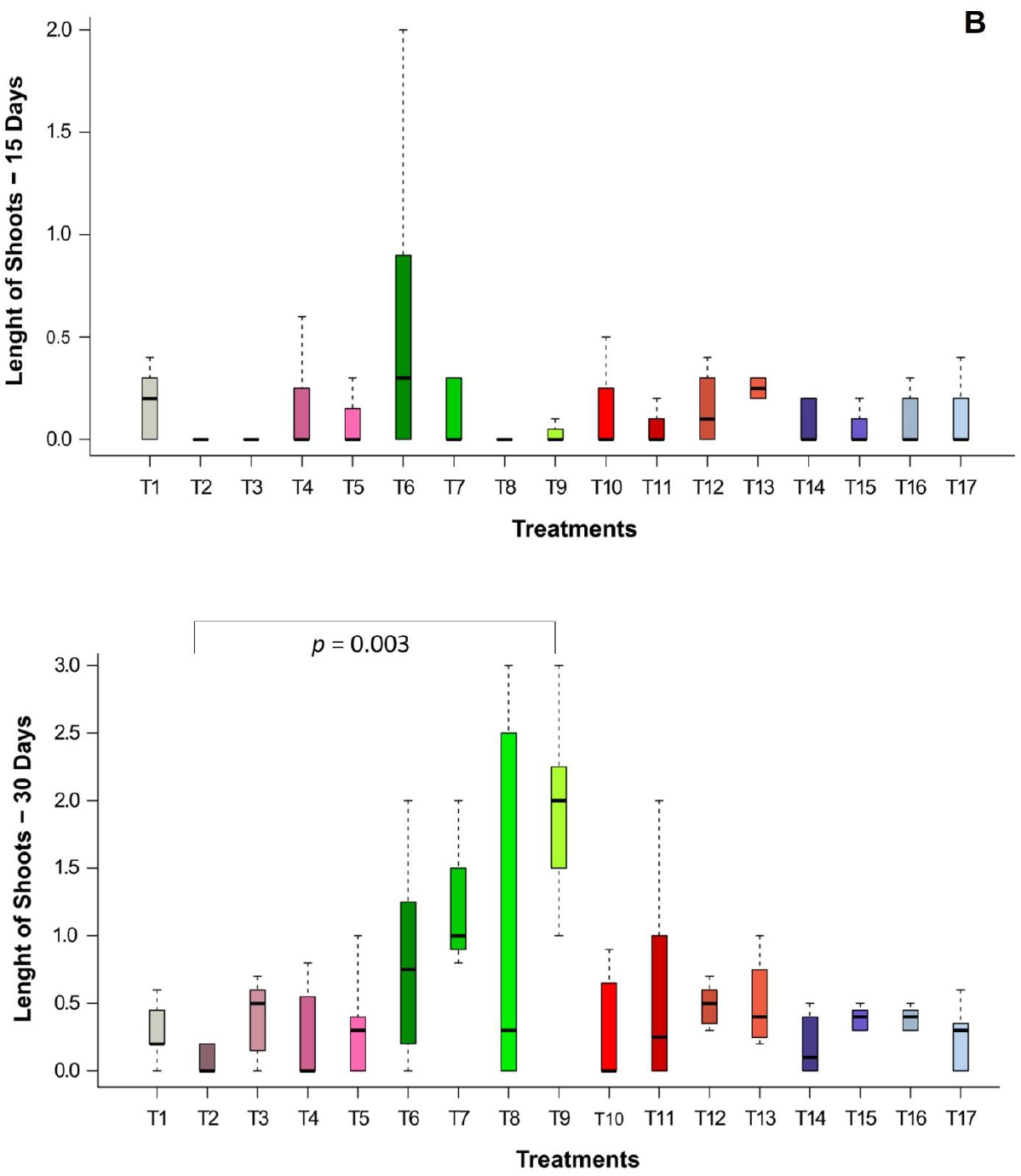
**a)** Number of shoots and **b)** length of shoots (15 and 30 days) on seedlings of a hybrid of *Phalaenopsis* Golden Peoker ‘BL’ HCC/AOS’ under the influence of isolated concentrations of BA and CIN and combined with NAA. Complete plants were submitted for treatments Control (T1), 1.0 mg BA (T2), 1.5 mg BA (T3), 2.0 mg BA (T4), 2.5 mg BA (T5), 1.0 mg BA + 0.5 mg NAA (T6), 1.5 mg BA + 0.5 mg NAA (T7), 2.0 mg BA + 0.5 mg NAA (T8), 2.5 mg BA + 0.5 mg NAA (T9), 1.0 mg KIN (T10), 1.5 mg KIN (T11), 2.0 mg KIN (T12), 2.5 mg KIN (T13), 1.0 mg KIN + 0.5 mg NAA (T14), 1.5 mg KIN + 0.5 mg NAA (T15), 2.0 mg KIN + 0.5 mg NAA (T16), 2.5 mg KIN + 0.5 mg NAA (T17). Data are presented as median (n =8). The Friedman test, followed by a *post-hoc* Bonferroni correction, was used for data analysis. The test result indicated that treatments differed significantly (*F*r = 32.436, *p* < 0.05).

Hyperhydricity and/or vitrification is a physiological state in which the plant has an abnormal water accumulation within the cells and tissues, resulting in a translucent and watery appearance [15-17]. Furthermore, hyperhydricity is a significant problem in the tissue culture industry since it may affect shoot multiplication and culture vigor and impede the successful transfer of micropropagated plants to *in vivo* conditions [17]. In this work, it was seen that the T13 (2.5 mg Kin), which showed signs of hyperhydricity, had no continuity in seedling formation and development, so acclimation was not possible.

The second highest median for number of shoots was the T6 (1.0 mg BA + 0.5 mg NAA), with 1.5 shoots per plant at 15 days. At 30 days, the second highest number of shoots was the T16 (2.0 mg KIN + 0.5 mg NAA), with 6.15 shoots per plant, and the T15 (1.5 mg KIN + 2.7 μM NAA), with 4.87 shoots per plant, followed by the T9 (2.5 mg BA + 0.5 μM NAA), with 4.25 shoots per plant (Fig. 2).

The work with orchids and their hybrids was developed based on the process of direct organogenesis as described by Gantait and Sinniah (18), using leaf segments [19], with meristematic apices of *Dendrobium nobile* and entire protocorms, or with the removed apex–thin cell layer–TCL of *Cymbidium* hybrids performed by Teixeira da Silva and Tanaka (20); Naing, Chung (21), working with *Coelogyne cristata* and Ng and Saleh (22), with the species *Paphiopedilum sp*.

Regarding shoot length, it was possible to see that, at 15 days of cultivation, the best treatment was the T6 (1.0 mg BA + 0.5 mg NAA) (Fig. 2). However, at 30 days, the one that stood out was the T9 (2.5 mg BA + 0.5 mg NAA) (Fig. 2).

Among the treatments, the T9 (2.5 mg BA + 0.5 mg NAA) showed the best development of shoots at the end of 30 days of cultivation in the organogenesis culture medium, as the median for shoot length was the highest. Morphologically, the shoots showed leaf expansion, subsequent plant elongation, and root formation, enabling the individualization of shoots and the efficient acclimatization of the plants at the end of the process (Figs. 1G and 1H).

Phytoregulators of the cytokinin class play a central role in the cell cycle influencing numerous development programs. From a historical viewpoint, they began having significance in studies because of a work conducted by Skoog and Miller (7), in which the authors found that phytoregulators of this class are substances that promote cell division.

The balance between cytokinins and auxins is relevant to controlling various aspects of cell differentiation and organogenesis in plant tissue culture (24) since the cytokinins break the apical dominance, influencing the multiplication rate, while the auxins help develop the culture (25). The BA growth regulator is a cytokinin widely used to stimulate the induction of shoots in ornamental plants (23).

Regarding the presence of root primordia in shoots, the best treatments were the T2 (1.0 mg BA), T3 (1.5 mg BA), T9 (2.5 mg BA + 0.5 mg NAA), T10 (1.0 mg Kin), and T17 (2.5 mg KIN + 0.5 mg NAA), all with the formation of root primordia at 15 days, remaining with these same treatments at 30 days of in vitro culture. The treatment with the lowest average for this variable was the T13 (2.5 mg Kin) during the 15^th^ and 30^th^ days of cultivation.

According to Pereira, Kasuya (23), roots formed in the in vitro cultivation of orchids may be related to the conventional type of morphology of orchid species, being epiphytic plants, and their physiology tends to have radicles and root primordia. However, for formed shoots, new roots are referred to the induction of phytoregulators, such as auxins, in the culture medium [24].

Using auxins favors the initiation of lateral root formation at specific sites within the basal meristem forming specific cell groups (XPP) with a particular molecular identity and sites of DR5 auxin response promoters able to respond to and initiate lateral root formation [25]. Auxin and cytokinin use can perform a favorable interaction for several developmental processes in plants, including lateral root development [25], having classical experiences that the balance between auxin and cytokinin becomes a key regulator in in vitro organogenesis [7].

Direct organogenesis developed with various species of orchids shows satisfactory results from the combination of cytokinin/auxin. As an example, working with *Dendrobium*, Talukder *et al*. (2003) obtained results in shoot proliferation, root formation, and shoot elongation from the combination of cytokinin/auxin (2.5 mg BA + 0.5 mg NAA).

During the acclimatization phase, the sphagnum substrate used for cultivation favored the plant development. This result was achieved as a result of the aeration of the roots and moisture conservation, mitigating the damage caused by the stress suffered by the environmental change [26]. The combination of 2.5 mg BA + 0.5 mg NAA (T9) in these cultivation conditions was the treatment that stood out in the three variables for the in vitro production of orchids of the *Phalaenopsis* Golden Peoker ‘BL’ HCC / AOS’ hybrid due to the formation of buds, which converted and differentiated into complete plants with the development of shoots and root primordia (Fig. 1H). This facilitated the acclimatization and survival of plants obtained from the organogenesis process (Fig. 1I).

## CONCLUSION

The asepsis and the in vitro germination of seeds from the direct organogenesis process proved efficient in producing the orchids of *Phalaenopsis* Golden Peoker ‘BL’ HCC/AOS’ hybrid. At 90 days of in vitro cultivation, the plants showed an adequate morphological structure to be used as a source of explants for inducing organogenesis, differentiation, and shoot development.

The T9 (11 μM BA + 2.7 μM NAA) was the most suitable for reproducing an efficient protocol since the treatment enabled the best development of the plants, with complete structures, satisfactory multiplication rate for the variable number of shoots, size, and development of the root system, significant aspects for reproducibility of a regeneration protocol.

## Acknowledgments

Thanks are due to DSc. Maurecilne Lemes da Silva Carvalho for supporting this study.

## Supplementary Material

Graph of the frequencies of observations *versus* treatments by analysis using the R software.

