## Supplementary figures and images for "The Auxin and Cytokinin Balance Influence the in Vitro Regeneration of *Phalaenopsis* Shoots (Orchidaceae)"

### Supplementary figure

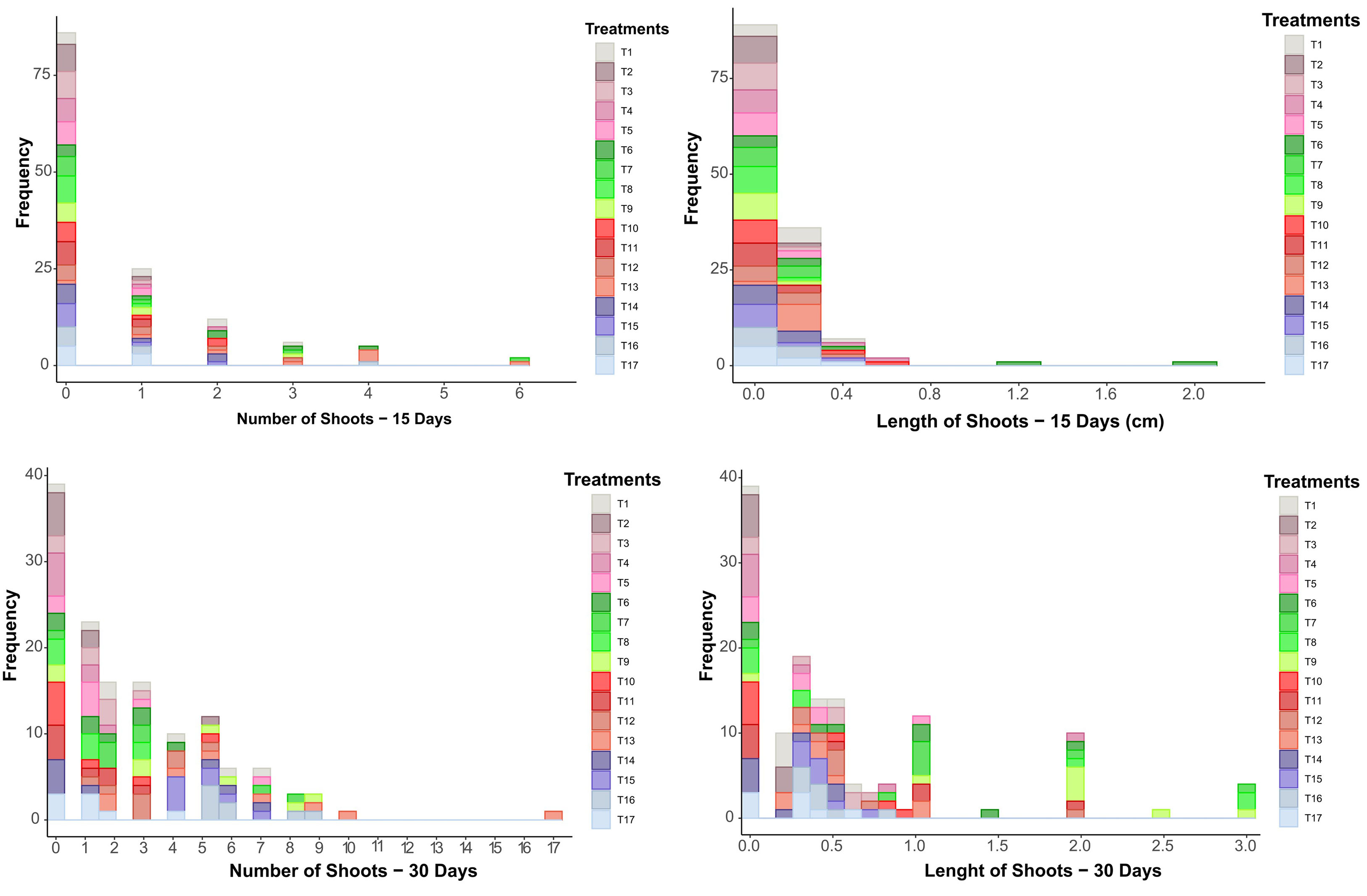
